# The Role of Sympathetic Innervation in Neonatal Muscle Growth and Neuromuscular Contractures

**DOI:** 10.1101/2023.06.20.545748

**Authors:** Mason T. Runkel, Albaraa Tarabishi, Kritton Shay-Winkler, Marianne E. Emmert, Qingnian Goh, Roger Cornwall

## Abstract

Neonatal brachial plexus injury (NBPI), a leading cause of pediatric upper limb paralysis, results in disabling and incurable muscle contractures that are driven by impaired longitudinal growth of denervated muscles. A rare form of NBPI, which maintains both afferent and sympathetic muscle innervation despite motor denervation, protects against contractures. We have previously ruled out a role for NRG/ErbB signaling, the predominant pathway governing antegrade afferent neuromuscular transmission, in modulating the formation of contractures. Our current study therefore investigated the contributions of sympathetic innervation of skeletal muscle in modulating NBPI-induced contractures. Through chemical sympathectomy and pharmacologic modification with a β_2_-adrenergic agonist, we discovered that sympathetic innervation alone is neither required nor sufficient to modulate contracture formation in neonatal mice. Despite this, sympathetic innervation plays an intriguing sex-specific role in mediating neonatal muscle growth, as the cross-sectional area (CSA) and volume of normally innervated male muscles were diminished by ablation of sympathetic neurons and increased by β-adrenergic stimulation. Intriguingly, the robust alterations in CSA occurred with minimal changes to normal longitudinal muscle growth as determined by sarcomere length. Instead, β-adrenergic stimulation exacerbated sarcomere overstretch in denervated male muscles, indicating potentially discrete regulation of muscle width and length. Future investigations into the mechanistic underpinnings of these distinct aspects of muscle growth are thus essential for improving clinical outcomes in patients affected by muscle disorders in which both length and width are affected.

## Introduction

Neonatal brachial plexus injury (NBPI) during childbirth is the most common cause of pediatric upper limb paralysis [1], and leads to the secondary formation of disabling and incurable muscle contractures, or joint stiffness [2, 3]. These contractures severely impede mobility of the affected limb, ultimately resulting in dysfunction and disability [4]. Previous work in animal and clinical studies has revealed that contractures are caused by impaired longitudinal growth of denervated muscles [5–12], which is driven by elevated levels of muscle proteasome activity [13–15]. It is thus imperative to decipher mechanistic links between denervation, muscle proteostasis, longitudinal muscle growth, and contractures in order to identify novel avenues for contracture prevention and treatment. The most common type of NBPI that causes contractures involves postganglionic nerve root injuries. This type of the injury occurs distal to the dorsal root ganglion and disrupts afferent (sensory), efferent (motor), and sympathetic innervation, leading to a completely denervated muscle [8]. In contrast, a rarer form of NBPI involves preganglionic nerve root avulsion, whereby nerve roots are injured proximal to the dorsal root ganglion [16]. Although this type of injury also disrupts efferent innervation, it preserves both afferent and sympathetic innervation, as it does not disrupt the connection between the muscle and the dorsal root ganglion containing the afferent neuron cell bodies or the sympathetic chain containing the sympathetic neuron cell bodies. Critical to the understanding of contractures is the fact that clinically, children with preganglionic NBPI do not develop severe contractures as seen in postganglionic NBPI [16, 17], despite identical motor paralysis as occurs in postganglionic NBPI. The potential protective effect of afferent and sympathetic innervation against contractures must therefore be investigated to decipher underlying molecular mechanism(s) by which neural input governs longitudinal muscle growth and contracture formation following neonatal denervation.

Both afferent and sympathetic innervation impact various aspects of skeletal muscle development and homeostasis. Afferent innervation in particular regulates muscle spindle formation and maintenance through the NRG/ErbB signaling pathway [18–21]. Utilizing mouse models of preganglionic and postganglionic NBPI, we previously reported that NRG/ErbB signaling is neither required nor sufficient for the modulation of contractures or longitudinal muscle growth [22]. While these findings do not directly rule out the role of afferent muscle innervation nor alternate/noncanonical pathways by which afferent innervation may be involved in contracture formation, they suggest that the difference in contractures between pre– and postganglionic NBPI may instead be modulated by sympathetic muscle innervation. Similar to afferent neurons, sympathetic neurons have been shown to directly innervate muscle spindles [23]. In addition, sympathetic input controls synapse maintenance and function at the neuromuscular junction, and regulates postsynaptic levels of membrane acetylcholine receptor [24]. These critical functions are exerted through β-adrenergic signaling [25], which serves a key role in regulating many aspects of muscle homeostasis, especially in diseased and dysfunctional muscles. Specifically, β-adrenergic signaling prevents muscle wasting in denervated muscles, and promotes regeneration after injury [26]. Importantly, treatment with β_2_-adrenergic agonists such as clenbuterol directly ameliorates denervation-induced atrophy through inhibition of muscle-specific proteasome activity [27], which we have shown to be integral to contracture formation. Despite this, the role of β-adrenergic signaling in longitudinal muscle growth remains to be explored.

In the current study, we dissected the role of sympathetic muscle innervation in modulating longitudinal muscle growth and contractures. Here, we specifically investigated whether the protective effect of preganglionic injury against contractures is conferred by sympathetic neural input, and whether restoration of β-adrenergic signaling prevents contractures after postganglionic injury. We surgically created different models of NBPI that preserved afferent/sympathetic muscle innervation or led to complete denervation, and proceeded to chemically ablate sympathetic neurons or pharmacologically stimulate β_2_-agonist activity, respectively. Our results showed that loss of sympathetic innervation alone does not impair longitudinal muscle growth or cause contractures in preganglionic injury, and that stimulation of β-adrenergic signaling alone is not sufficient to rescue longitudinal muscle growth or prevent contractures in postganglionic injury. However, alterations in this neuronal pathway impacted growth of normally innervated muscles in male mice. Furthermore, the sex-specific increase in cross-sectional area of normally innervated muscles with pharmacologic β_2_-agonist treatment was accompanied by worsened longitudinal growth impairment of denervated muscles. Our collective findings thus rule out a mechanistic role of sympathetic innervation in governing contracture formation and eliminate β_2_-agonists as a pharmacologic target for modulating contractures. However, our investigation also identifies a sex-specific role of the sympathetic network in governing neonatal muscle growth and reveals an intriguing divergence in the regulation of muscle width and length.

## Results

### Loss of sympathetic innervation does not elicit contractures

To investigate a possible isolated role of sympathetic innervation in longitudinal muscle growth and contractures, we first asked whether sympathetic denervation alone was sufficient to induce contractures without NBPI. Following surgical excision of the cervical sympathetic ganglia in neonatal rats (**Fig. 1A**), there were no alterations in elbow joint range of motion (**Figs. 1B–D**) or sarcomere length of the brachialis muscles (**Figs. 1E,F**). Curiously, we observed a selective inhibition of sympathetic axons along the sympathetic chain of the MCN that branches into the biceps muscle, but not the chain branching into the sensory portion of the lateral antebrachial cutaneous nerve (LABCN) (**Fig. 1 – suppl. fig 1A**). We therefore treated neonatal mice with the neurotoxin 6-hydroxydopamine (6-OHDA) (**Fig. 2A**), which specifically destroys all sympathetic neurons in the sympathetic ganglia [28]. This robust means of chemical sympathectomy resulted in the complete destruction of sympathetic axons along both the biceps and LABCN branches of the MCN (**Fig. 2 – suppl. fig 1A)**. Four weeks post treatment, there similarly were no changes in passive elbow extension (**Figs. 2B,C**), or longitudinal growth of the brachialis (**Figs. 2D,E**). These overall findings confirm that in a non-NBPI environment, loss of sympathetic innervation does not mediate contracture formation.

**Figure 1:**
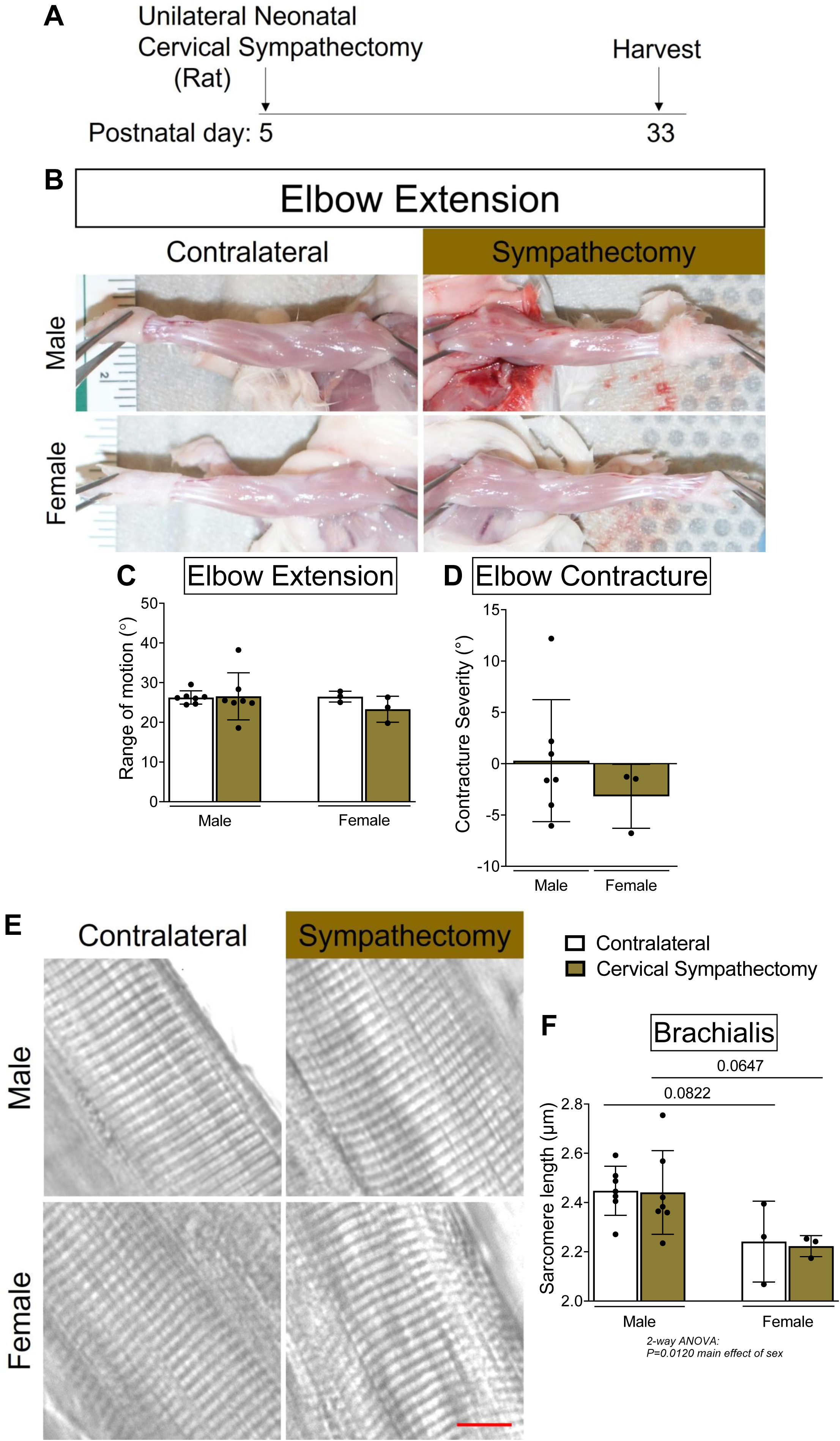
Surgical sympathectomy does not induce contractures. (**A**) Schematic depicting cervical sympathectomy in neonatal rats (P5) and assessment of joint motion at 4 weeks post-surgery (P33). (**B**) Representative images of sympathectomized and contralateral forelimbs, and quantitation of both (**C**) elbow joint range of motion and (**D**) corresponding elbow flexion contracture severity revealed that loss of sympathetic innervation alone does not induce contractures in rats. (**E**) Representative DIC images of sarcomeres, and (**F**) quantitation of sarcomere length further showed that cervical sympathectomy alone fails to perturb functional length in sympathectomized rat muscles. In (**D**), contracture severity is calculated as the difference in passive elbow extension and shoulder rotation between the sympathectomized side and the contralateral side. Data are presented as mean ± SD, n = 9 independent rats. Statistical analyses: (**C**), (**F**) two-way ANOVA for sex and surgery (repeated measures between forelimbs) with a Bonferroni correction for multiple comparisons, (**D**) unpaired two-tailed Student’s t-tests. Scale bar (**E**): 10 µm.

**Figure 2:**
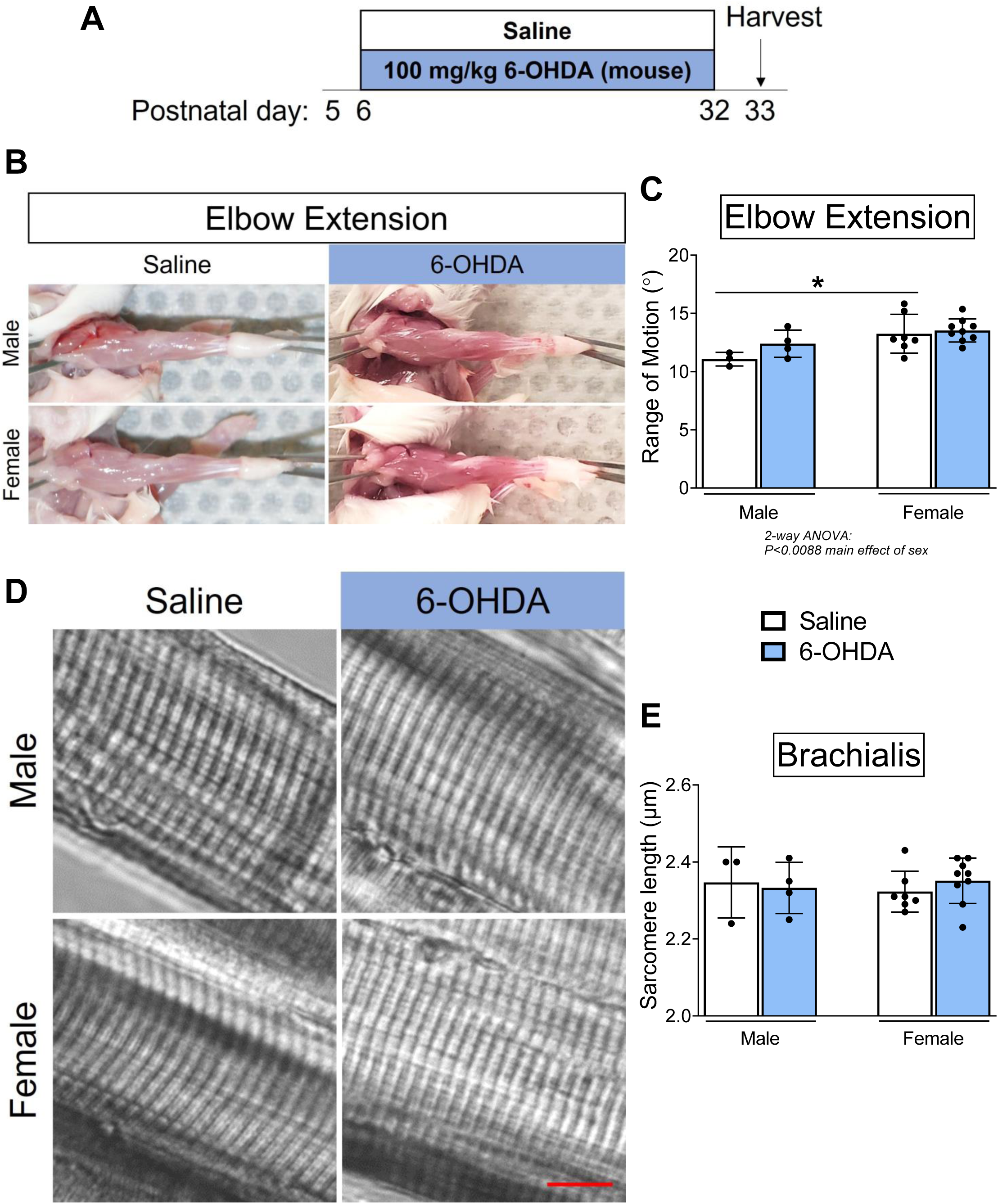
Chemical sympathectomy does not induce contractures. (**A**) Schematic depicting saline or 100 mg/kg 6-OHDA treatment to chemically ablate sympathetic neurons in mice during neonatal muscle development (P6 – P32), and assessment of joint motion at P33. (**B**) Representative images of forelimbs and (**C**) quantitative analyses of elbow extension showed that absence of sympathetic neurons does not reduce elbow joint range of motion in mice. (**D**) Representative DIC images of sarcomeres, and (**E**) quantitation of sarcomere length further revealed chemical sympathectomy alone does not alter the functional length of sympathectomized mouse muscles. Data are presented as mean ± SD, n = 3-9 independent mice per group. Statistical analyses: (**C**), (**E**) two-way ANOVA for sex and treatment with a Bonferroni correction for multiple comparisons. **\***p<0.05. Scale bar (**D**): 10 µm.

We next utilized chemical sympathectomy with 6-OHDA to interrogate the role of sympathetic innervation in modulating contractures after neonatal denervation. If the protective effect of preganglionic injury against contractures in paralyzed muscles is governed by sympathetic innervation, we would then expect that ablation of sympathetic axons would lead to the formation of contractures after preganglionic NBPI, which does not cause contractures despite motor paralysis of affected muscles [8, 16, 17]. Following unilateral preganglionic NBPI surgery (**Fig. 3A**), neonatal P5 mice were subjected to 4 weeks of 6-OHDA to ablate sympathetic neurons (**Fig. 3B**). While preganglionic injury alone restricted elbow extension (**Fig. 3D**) by ∼10° in male mice, this is within the 10° threshold for non-clinically relevant differences. This subtle alteration in range of motion after preganglionic NBPI corresponded to a mild sarcomere overstretch of 0.02 µm in brachialis muscles (**Fig. 3G**), similar to our previously reported observation [22]. Importantly, loss of sympathetic innervation did not further restrict joint range of motion, nor induce the formation of contractures at the elbow joint in either male or female mice (**Figs. 3C–E**). Furthermore, chemical sympathectomy did not reduce functional muscle length in denervated forelimbs, as brachialis sarcomere length was not perturbed by 6-OHDA treatment in pre-ganglionic NBPI or contralateral control limbs (**Figs. 3F,G**). Thus, the absence of contractures and sarcomere overstretch with 6-OHDA treatment after preganglionic NBPI reveals that the protective effect of preganglionic injury is not conferred by sympathetic innervation alone.

**Figure 3:**
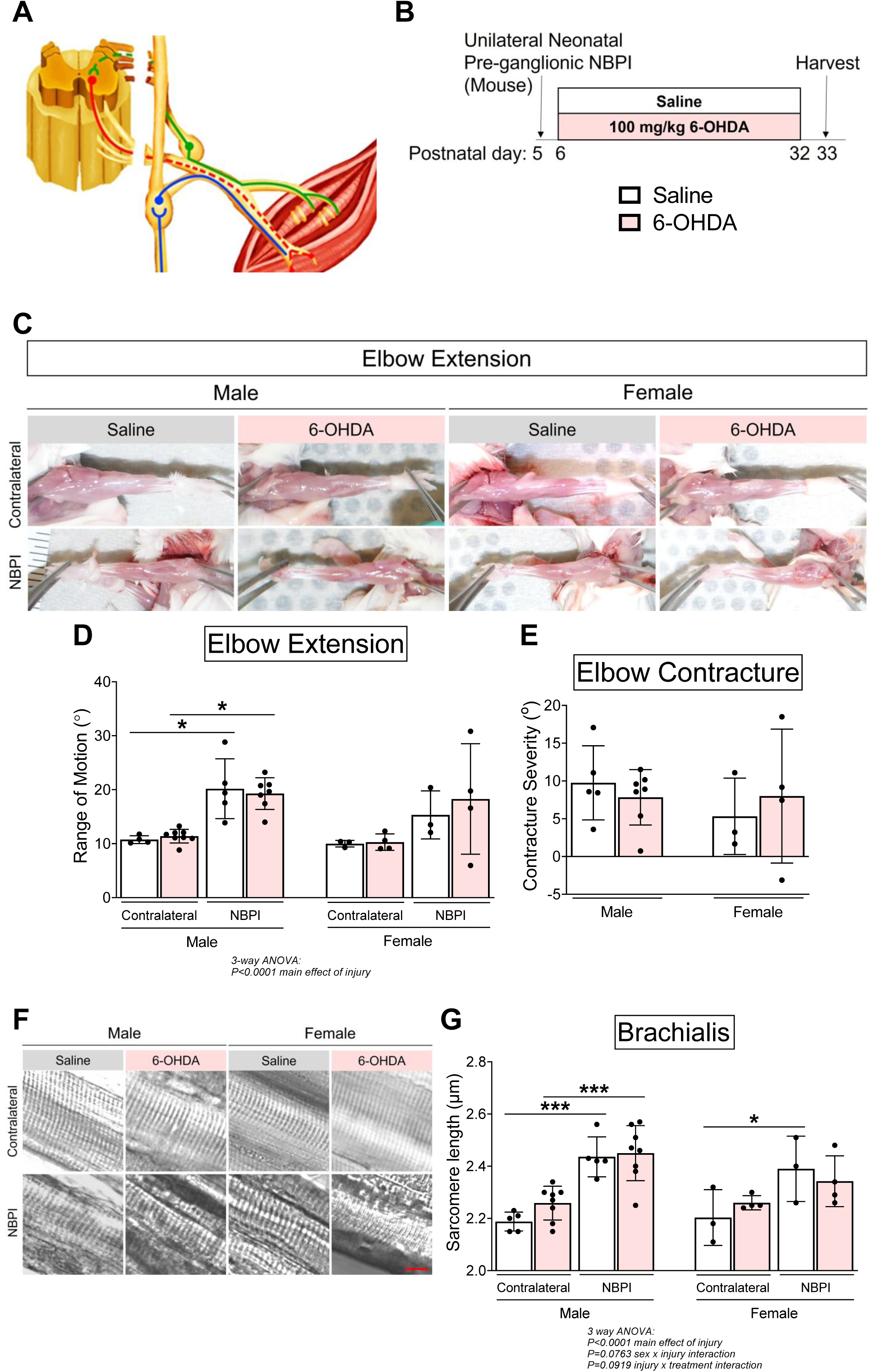
Cervical sympathectomy does not induce contractures after preganglionic injury. (**A**) Schematic diagram of preganglionic NBPI, with efferent (red), afferent (green), and sympathetic (blue) neurons depicted. Dashed lines indicate Wallerian degeneration of axons (denervation). (**B**) Schematic depicting saline or 6-OHDA treatment during neonatal muscle development (P8 – P32), and assessment of joint motion at 4 weeks following a mouse model of preganglionic NBPI. (**C**) Representative images of denervated (NBPI) and contralateral forelimbs, and quantitation of both (**D**) elbow joint range of motion and (**E**) contracture severity revealed that loss of sympathetic innervation does not promote contracture formation. (**F**) Representative DIC images of sarcomeres, and (**G**) quantitation of sarcomere length also showed that while preganglionic NBPI leads to sarcomere elongation, chemical sympathectomy does not further impair functional length in denervated (NBPI) muscles. In (**E**), elbow flexion contracture severity is calculated as the difference in passive elbow extension and between the NBPI and contralateral sides. Data are presented as mean ± SD, n = 3-8 independent mice per group. Statistical analyses: (**D**), (**G**) three-way analysis of variance (ANOVA) for sex, treatment, and denervation (repeated measures between forelimbs) with a Bonferroni correction for multiple comparisons, (**E**) two-way ANOVA for sex and treatment with a Bonferroni correction for multiple comparisons. **\***p<0.05, **\*\*\***p<0.001. Scale bar (**F**): 10 µm.

### Sex-specific requirement of sympathetic neurons for neonatal muscle growth

We subsequently extended our investigation on the role of sympathetic innervation to different aspects of musculoskeletal development. Four weeks of chemical sympathectomy reduced cross-sectional and volumetric growth of normally innervated brachialis muscles in male neonatal mice by ∼16% and ∼18%, respectively (**Figs. 4A–D**). Despite this growth deficit, no differences were observed in CSA and overall volume of denervated muscles across the sexes with preganglionic injury. Instead, the size attenuation in innervated muscles with 6-OHDA was accompanied by a reduction in terminal body weight (**Fig. 4 – suppl. fig 1A**), without a corresponding impairment in skeletal growth as measured by humerus length (**Fig. 4 – suppl. fig 1B**). These results indicate that body weight losses are attributed mainly to deficient neonatal muscle growth.

**Figure 4:**
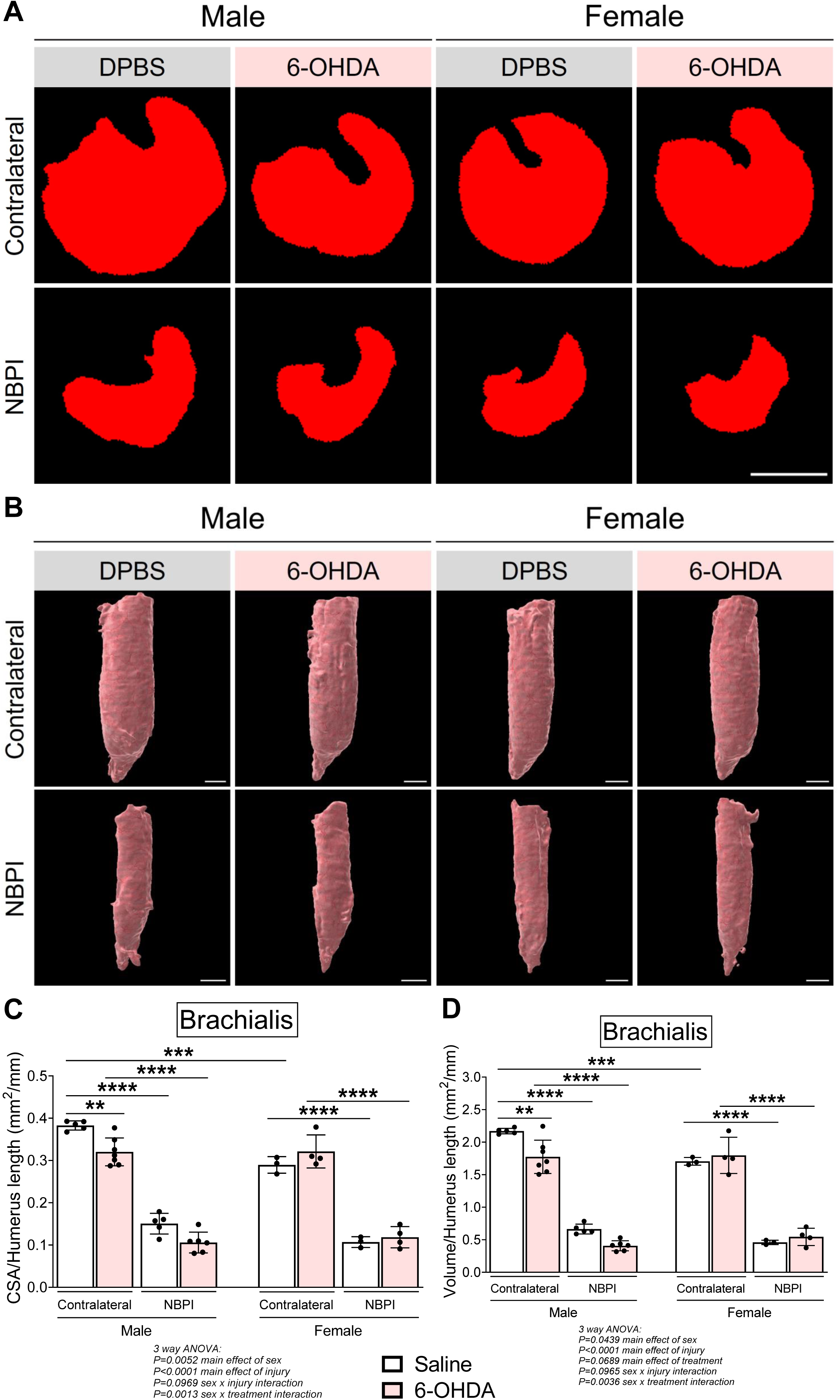
Cervical sympathectomy impairs neonatal muscle growth in a sex-specific manner. Representative micro-CT images in (**A**) transverse and (**B**) 3-dimensional views revealed smaller brachialis muscles in normal innervated forelimbs of male mice with chemical sympathectomy. (**C**) Analyses of brachialis CSA and (**D**) volume further confirmed that neonatal growth of normally innervated brachialis muscles in contralateral forelimbs is blunted with ablation of sympathetic axons only in male mice. Data are presented as mean ± SD, n = 3-7 independent mice per group. Statistical analyses: (**C**), (**D**) three-way analysis of variance (ANOVA) for sex, treatment, and denervation (repeated measures between forelimbs) with a Bonferroni correction for multiple comparisons. **p<0.01, ***p<0.001, ****p<0.0001. Scale bars (**A**), (**B**): 1000 µm.

### β-adrenergic stimulation does not prevent contractures

Having eliminated the role of sympathetic innervation in a non-contracture NBPI model, we proceeded to dissect its contributions in contracture formation after postganglionic injury which does cause contractures. Here, we tested the hypothesis that in an injury model which causes contractures, increased β-adrenergic signaling would be sufficient to prevent contractures. Following unilateral postganglionic NBPI surgery (**Figs. 5A, 6A**), neonatal P5 mice and rats were treated daily for 4 weeks with clenbuterol, a β_2_-adrenergic receptor agonist (**Figs. 5B, 6B**). β-adrenergic stimulation failed to improve passive elbow extension (**Figs. 5C,D**; **Figs. 6C,D**), resulting in an ineffective prevention of elbow flexion contractures (**Figs. 5E, 6E**) in both male and female mice, as well as male and female rats. Furthermore, β-adrenergic stimulation did not preserve longitudinal muscle growth, as sarcomere overstretch was further exacerbated solely in denervated brachialis muscles of male mice and rats with clenbuterol treatment (**Figs. 5F,G**; **Figs. 6F,G**). This additional reduction in functional muscle length did not worsen elbow contracture severity, supporting our prior observations of a threshold-dependent role of sarcomere length in contracture formation [22]. These findings thereby indicate that increased β-adrenergic signaling alone is insufficient to correct deficits in muscle length and prevent the formation of contractures after postganglionic injury.

**Figure 5:**
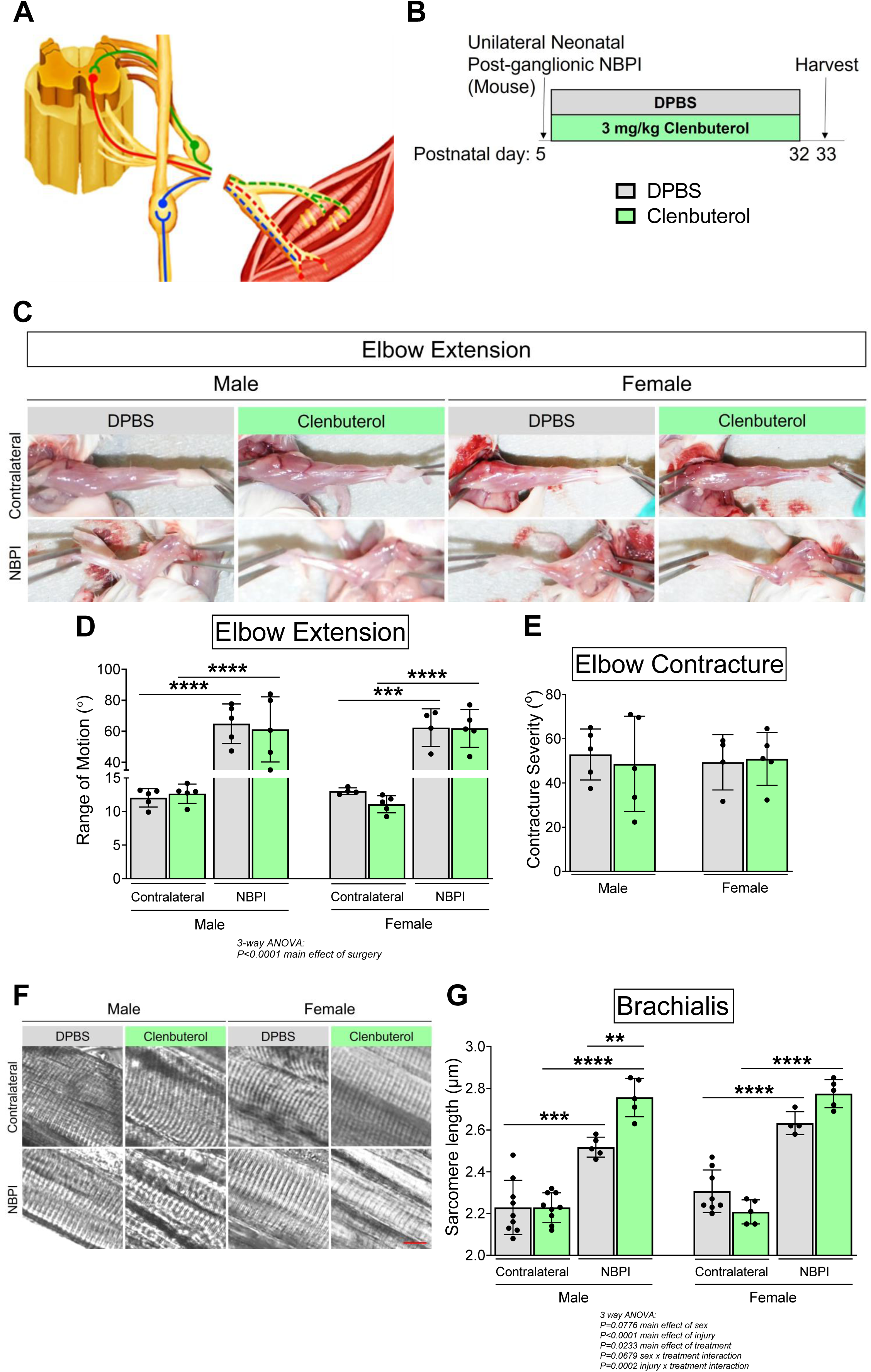
β-adrenergic stimulation does not prevent contractures after postganglionic injury in mice. (**A**) Schematic of postganglionic NBPI, with efferent (red), afferent (green), and sympathetic (blue) neurons depicted. Dashed lines indicate Wallerian degeneration of axons (denervation). (**B**) Schematic depicting Dulbecco’s phosphate-buffered saline (DPBS) or 3 mg/kg Clenbuterol from P8 – P32, and assessment of joint motion at 4 weeks following a mouse model of postganglionic NBPI. (**C**) Representative images of denervated (NBPI) and contralateral forelimbs, and quantitative analyses of both (**D**) elbow joint range of motion and (**E**) contracture severity revealed that β-adrenergic activation does not prevent contracture formation in mice. n = 4-5 independent mice per group. (**F**) Representative DIC images of sarcomeres further showed that clenbuterol treatment does not reduce sarcomere overstretch in denervated brachialis muscles of mice after postganglionic NBPI. Instead, quantitation of (**G**) absolute sarcomere length revealed additional overstretch with β-adrenergic stimulation following postganglionic injury in male mice. n = 4-9 independent mice per group. In (**E**), elbow flexion contracture severity is calculated as the difference in passive elbow extension between the NBPI and contralateral sides. Data are presented as mean ± SD. Statistical analyses: (**D**), (**G**) three-way analysis of variance (ANOVA) for sex, treatment, and denervation (repeated measures between forelimbs) with a Bonferroni correction for multiple comparisons, (**E**) two-way ANOVA for sex and treatment with a Bonferroni correction for multiple comparisons. **p<0.01, ***p<0.001, ****p<0.0001. Scale bar (**F**): 10 µm.

**Figure 6:**
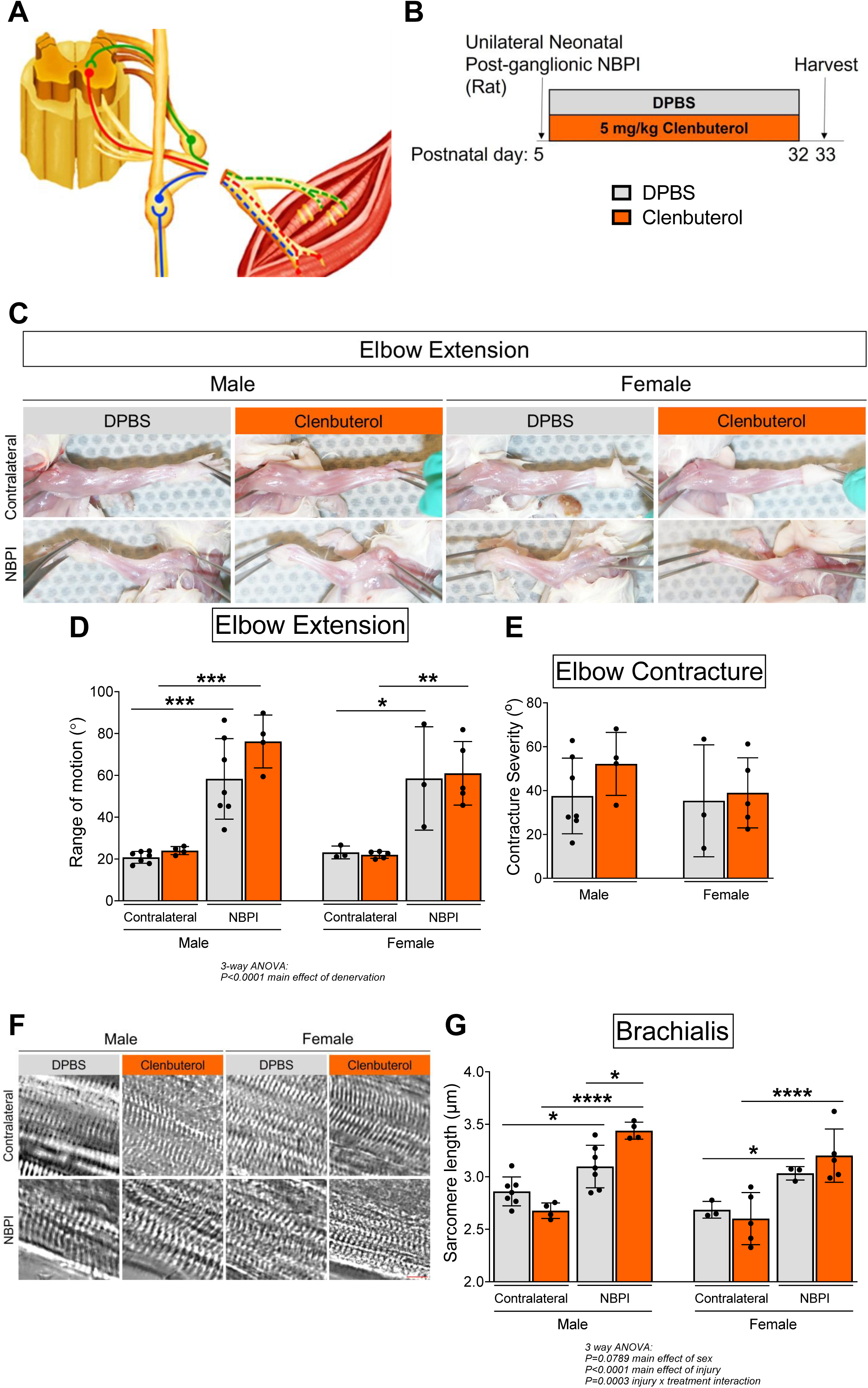
β-adrenergic stimulation does not prevent contractures after postganglionic injury in rats. (**A**) Schematic of postganglionic NBPI, with efferent (red), afferent (green), and sympathetic (blue) neurons depicted. Dashed lines indicate Wallerian degeneration of axons (denervation). (**B**) Schematic depicting Dulbecco’s phosphate-buffered saline (DPBS) or 5 mg/kg Clenbuterol from P8 – P32, and assessment of joint motion at 4 weeks following a rat model of postganglionic NBPI. (**C**) Representative images of denervated (NBPI) and contralateral forelimbs, and quantitative analyses of both (**D**) elbow joint range of motion and (**E**) contracture severity revealed that β-adrenergic activation does not prevent contracture formation in rats. (**F**) Representative DIC images of sarcomeres further showed that clenbuterol treatment does not reduce sarcomere overstretch in denervated brachialis muscles of rats after postganglionic NBPI. Instead, quantitation of (**G**) absolute sarcomere length revealed additional overstretch with β-adrenergic stimulation following postganglionic injury in male rats. In (**E**), elbow flexion contracture severity is calculated as the difference in passive elbow extension between the NBPI and contralateral sides. Data are presented as mean ± SD, n = 3-7 independent rats per group. Statistical analyses: (**D**), (**G**) three-way analysis of variance (ANOVA) for sex, treatment, and denervation (repeated measures between forelimbs) with a Bonferroni correction for multiple comparisons, (**E**) two-way ANOVA for sex and treatment with a Bonferroni correction for multiple comparisons. *p<0.05, **p<0.01, ***p<0.001, ****p<0.0001. Scale bar (**F**): 10 µm.

To complement our pharmacologic approach, we created a surgical rat model of neonatal deafferentation and deefferentation, which preserves sympathetic innervation to the muscle (**Fig. 7A**). The presence of elbow flexion contractures in both male and female rats suggests that in an otherwise denervated muscle, sympathetic innervation alone is not sufficient to prevent contracture formation (**Figs. 7B–D**). Likewise, the maintenance of sympathetic innervation alone in an otherwise denervated muscle does not prevent sarcomere overstretch (**Figs. 7E–F**). These findings thus recapitulate our above discovery that restoration of ꞵ-adrenergic signaling with Clenbuterol does not preserve longitudinal muscle growth nor prevent contractures in completely denervated muscles.

**Figure 7:**
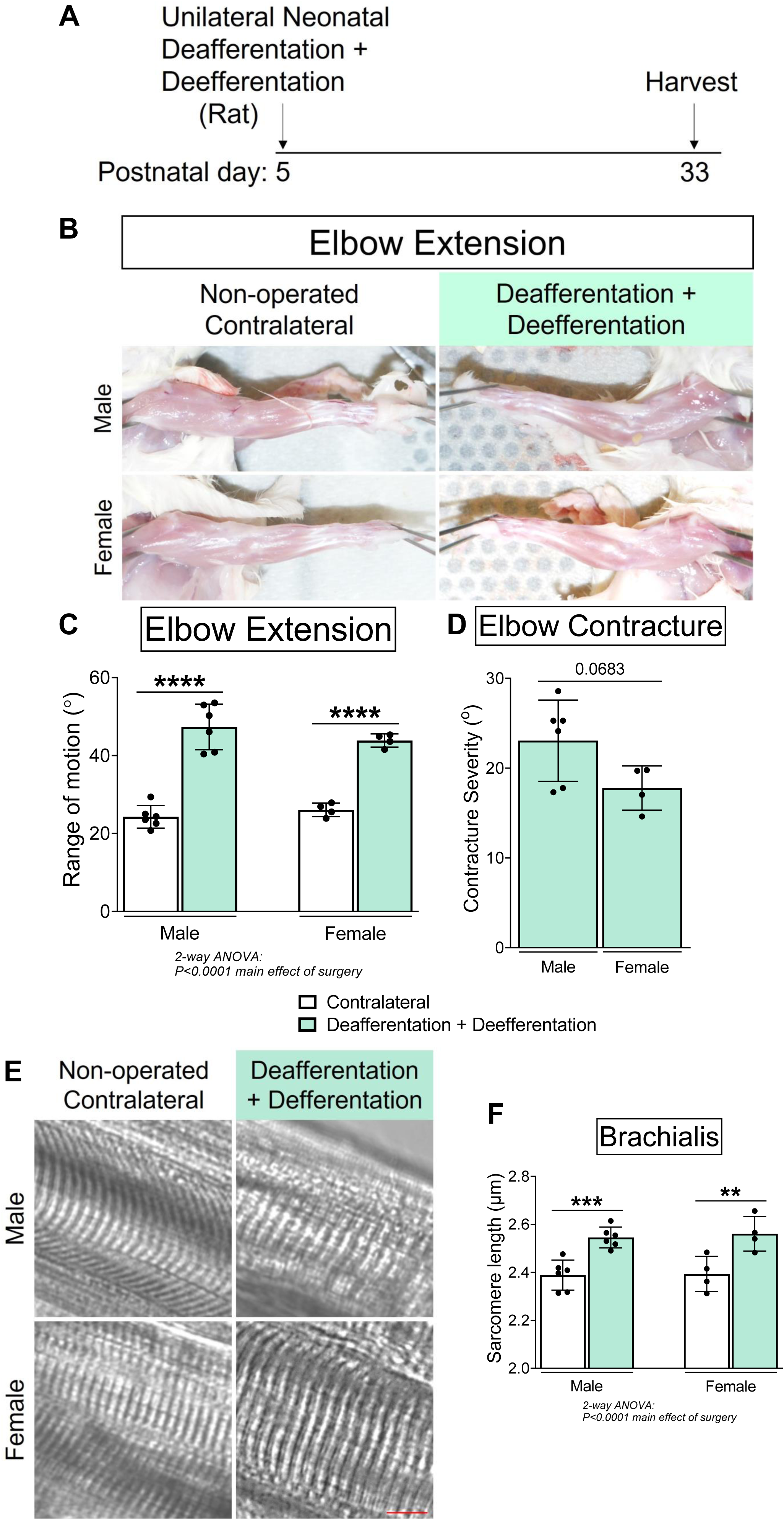
**Sympathetic muscle innervation is insufficient to prevent contractures**. (**A**) Schematic depicting combined deafferentation and deefferentation in neonatal rats (P5), and subsequent assessment of joint motion at 4 weeks post-surgery (P33). (**B**) Representative images of denervated and contralateral forelimbs, and quantitation of both (**C**) elbow joint range of motion and (**D**) contracture severity revealed that preservation of sympathetic muscle innervation alone is insufficient to prevent contractures in an otherwise denervated muscle. In (**D**) elbow flexion contracture severity is calculated as the difference in passive elbow extension between the surgical and contralateral sides. Clinically relevant contractures are defined as a ≥10° difference between sides. (**E**) Representative DIC images of sarcomeres, and (**F**) quantitation of sarcomere length revealed the presence of sarcomere overstretch in denervated muscles, indicating that sympathetic innervation alone does not restore longitudinal muscle growth. Data are presented as mean ± SD, n = 10 independent rats. Statistical analyses: (**C**), (**F**) two-way ANOVA for sex and surgery (repeated measures between forelimbs) with a Bonferroni correction for multiple comparisons, (**D**) unpaired two-tailed Student’s t-tests. **p<0.01, ***p<0.001, ****p<0.0001. Scale bar (**E**): 10 µm.

### β-adrenergic signaling promotes sex-specific neonatal muscle growth

Despite impairing functional length in denervated muscles, β-adrenergic stimulation with clenbuterol increased CSA and volume of normally innervated muscles solely in neonatal male mice by ∼20% and ∼17%, respectively (**Figs. 8A–D**). This sex dimorphism in the adrenergic effect on muscle CSA is not species-specific, as CSA (but not volume) of innervated muscles also increased by ∼27% solely in neonatal male rats after clenbuterol treatment (**Figs. 9A–D**). Given that the total volume of a muscle is a function of both its CSA and length, our results showed that the sex-dependent gains in muscle CSA with clenbuterol in both species are not accompanied by concurrent gains in muscle length. Rather, clenbuterol treatment preferentially increased muscle cross-sectional growth at the expense of longitudinal muscle growth. These collective findings reveal that β-adrenergic signaling regulates CSA and length in a divergent manner, indicating that muscle width and length are potentially governed through distinct mechanisms. Despite the sex dimorphism in innervated muscles, clenbuterol did not attenuate denervation-induced muscle atrophy in either male or female mice or rats (**Figs. 8A–D**; **Figs. 9A–D**). Instead, clenbuterol impaired humerus length of both forelimbs solely in female mice without reducing terminal body weight (**Fig. 8 – suppl. figs 1A,B**), suggesting a sex-specific attenuation of skeletal growth.

**Figure 8:**
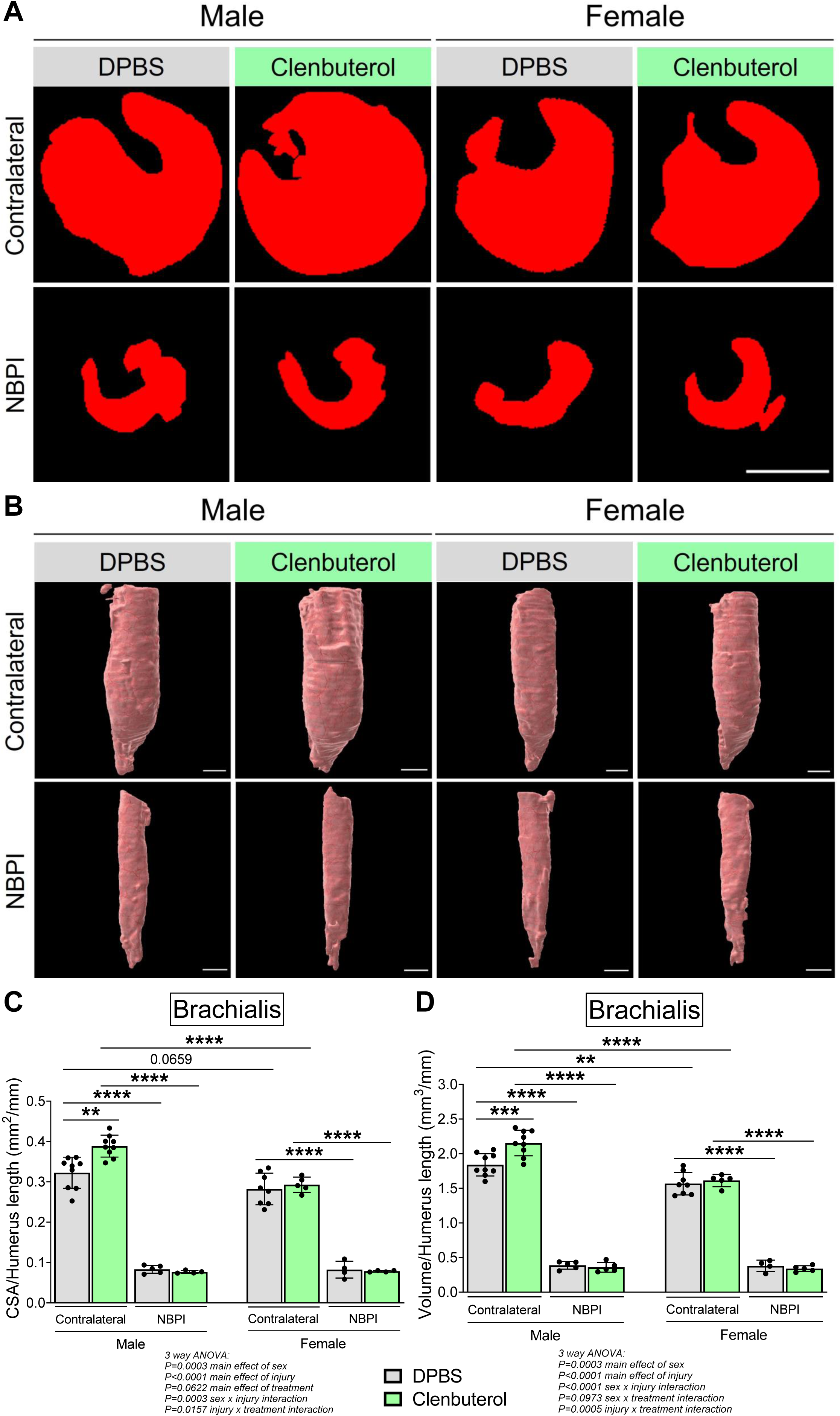
β-adrenergic stimulation enhances neonatal muscle growth in a sex-specific manner in mice. Representative micro-CT images in (**A**) transverse and (**B**) 3-dimensional views revealed larger brachialis muscles in normal innervated forelimbs of male mice with clenbuterol treatment. (**C**) Analyses of brachialis CSA and (**D**) volume further confirmed that neonatal growth of normally innervated brachialis muscles in contralateral forelimbs is increased with β-adrenergic activation only in male mice. Data are presented as mean ± SD, n = 4-9 independent mice per group. Statistical analyses: (**C**), (**D**) three-way analysis of variance (ANOVA) for sex, treatment, and denervation (repeated measures between forelimbs) with a Bonferroni correction for multiple comparisons. **p<0.01, ***p<0.001, ****p<0.0001. Scale bars (**A**), (**B**): 1000 µm.

**Figure 9:**
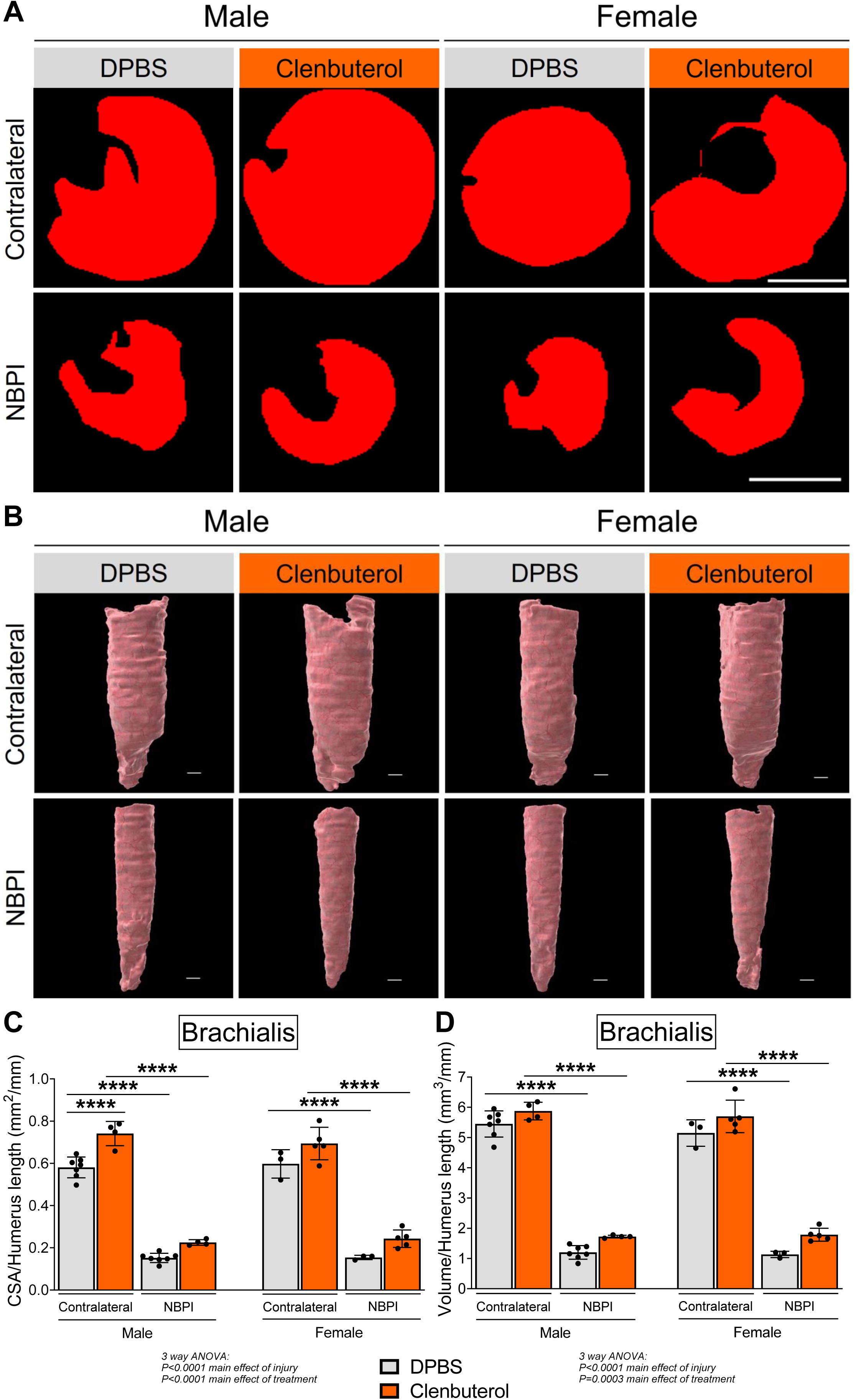
β-adrenergic stimulation enhances neonatal muscle growth in a sex-specific manner in rats. Representative micro-CT images (**A**) transverse and (**B**) 3-dimensional views revealed larger cross-sectional area of brachialis muscles in normal innervated forelimbs of male rats with clenbuterol treatment. Morphometric analyses of (**C**) cross-sectional area and (**D**) total volume revealed β-adrenergic activation increased cross-sectional (but not volumetric) growth of normally innervated brachialis muscles only in neonatal male rats. Data are presented as mean ± SD, n = 3-7 independent rats per group. Statistical analyses: (**C**), (**D**) three-way analysis of variance (ANOVA) for sex, treatment, and denervation (repeated measures between forelimbs) with a Bonferroni correction for multiple comparisons. ****p<0.0001. Scale bar (**A**), (**B**): 1000 µm.

## Discussion

Current strategies for restoring physical function to neonatally denervated limbs, including reconstructive surgery and manual rehabilitation, are unfortunately limited in their overall effectiveness. As existing contracture treatments have traditionally focused on palliative mechanical solutions instead of addressing the underlying pathophysiology of contracture formation, they may further impair function by weakening existing abnormal muscles [29–31]. The dearth in our understanding of contracture pathophysiology therefore represents a significant limitation to restoring muscle function and joint mobility to denervated limbs in children affected by neonatal brachial plexus injury. Important mechanistic insights into contracture development can be gained from preganglionic nerve root injuries, which preserve both afferent and sympathetic neural input to the muscle. Clinically, children with preganglionic NBPI do not develop severe contractures, despite incurring identical motor paralysis as occurs in postganglionic NBPI, which causes complete denervation and elicits severe and progressive contractures [16, 17]. This difference in contracture phenotype between injury modes challenges the long-held notion that contractures inevitably result from muscle paralysis [4, 32], and instead suggests that unique features of preganglionic NBPI protect against contractures even in paralyzed muscle [32]. Specifically, the preservation of afferent and/or sympathetic muscle innervation in preganglionic injuries could potentially sustain the muscle’s longitudinal growth, thereby protecting it against contractures despite motor paralysis. Rigorous interrogation of the contributions of both types of neuronal innervation in modulating contracture formation is critical to elucidate precise mechanistic links between denervation, muscle length, and contractures, and to enhance our understanding of the underlying mechanisms by which neural input impacts longitudinal muscle growth.

Utilizing validated mouse models that recapitulate the contracture phenotypes of both types of NBPI, we previously showed that NRG/ErbB signaling, the primary pathway governing antegrade afferent neuromuscular transmission, does not modulate the formation of contractures [22]. In this current study, we investigated whether sympathetic muscle innervation confers the protective effect of preganglionic injury against contractures, and whether recapitulation of β-adrenergic signaling, a key pathway governing sympathetic neural input, prevents contractures in completely denervated muscles after postganglionic injury. Our findings revealed that chemical sympathectomy with a neurotoxin, 6-OHDA, did not induce the formation of contractures following preganglionic NBPI. In addition, we found that stimulation with a β_2_-agonist, clenbuterol, failed to prevent postganglionic NBPI-induced contractures. Moreover, the preservation of sympathetic muscle innervation alone is insufficient to prevent contractures in a denervation model that disrupts both afferent and efferent innervation. These results therefore rule out a role for sympathetic muscle innervation and its β-adrenergic pathway in the modulation of contractures. By ruling out isolated roles for sympathetic innervation, β-adrenergic signaling, and NRG/ErbB signaling, we move closer towards deciphering underlying mechanisms by which innervation governs longitudinal muscle growth and contracture formation. Our current findings thus represent an important step forward by narrowing the search for pharmacologic targets for contracture prevention and treatment.

Our collective results have thus far demonstrated that the protective effect of preganglionic NBPI is neither conferred through NRG/ErbB signaling nor sympathetic innervation alone. It is important to note that we have not mechanistically ruled out other afferent contributions in contracture modulation. To this end, preliminary observations from neonatal rat models of deafferentation reveal that surgical excision of the dorsal root ganglion, which contains the afferent neuron cell bodies, does not cause contractures unless accompanied by motor paralysis (**data not shown**). More rigorous investigations are needed to fully decipher whether afferent innervation alone confers the protective effect of preganglionic injury against contractures. Beyond afferent neural input, we must also consider alternative mechanisms by which preganglionic injury protects against contractures. As we previously discussed, there may be alternate molecular pathways besides NRG/ErbB signaling that mediate the cross-talk between afferent neurons and skeletal muscle [22]. Another potential mechanism involves neurotrophins, a family of secreted growth factors that mediate the survival, function, and development of different sub-populations of sensory neuron in the dorsal root ganglia [33]. In particular, neurotrophin-3 (NT3) signaling through its high-affinity trkC receptors is essential for both the survival and differentiation of muscle spindle afferents [34, 35]. Lastly, our prior investigations into the protective effect of afferent innervation have focused primarily on the muscle spindles, which we have shown to be preserved in preganglionic injury, but completely degenerated following postganglionic NBPI [8]. However, we must rigorously explore the mechanistic roles of other mechanoreceptors that remain innervated after preganglionic NBPI. Indeed, our preliminary observations reveal preganglionic injury partially preserves the Golgi tendon organ whereas postganglionic NBPI disrupts it (**data not shown**), indicating a putative role for this receptor in protecting against contractures.

Additionally, while we can conclusively rule out the necessity of sympathetic innervation in protecting against contractures following preganglionic NBPI, we must continue to explore non-β-adrenergic mechanisms in preventing postganglionic NBPI-induced contractures. Sympathetic neurons can produce the pleiotropic cytokine, Interleukin 6 (IL-6), and respond to it in an autocrine/paracrine manner [36]. IL-6 myokine signaling in skeletal muscles serves paradoxical roles as it facilitates muscle stem (satellite) cell fusion, hypertrophy, and energy metabolism, while also promoting atrophy and wasting [37]. Indeed, pharmacological targeting of IL-6 and the IL-6/JAK/STAT3 pathway led to a rescue in denervated muscle atrophy [38]. Hence, this pathway is a potential signaling mechanism by which neural input regulates longitudinal muscle growth. Lastly, our studies up to this point have focused on the individual contributions of afferent and sympathetic innervation. We must thus continue to investigate the potential combined roles of both afferent and sympathetic neural input to the paralyzed muscle in protecting against contractures.

Besides investigating these neuronal pathways, we must also consider the role of other neural/glial cells, such as Schwann cells, which play a vital role in supporting axonal regeneration and the remyelinating of axons in denervated muscles [39]. In addition to facilitating nerve integrity and conduction, Schwann cells are required for the formation and maintenance of the neuromuscular junction, as their targeted ablation results in fewer and smaller acetylcholine receptor (AChR) clusters, as well as diminished miniature endplate potential amplitude and frequency [40]. Bulk and single cell gene expression analyses further revealed that the related muscle glial cells activate a neurotrophic signaling pathway that promotes neurite outgrowth and AChR clustering during nerve injury [41]. Through these mechanisms, Schwann cells and skeletal muscle-resident glial cells maintain muscle homeostasis in response to denervation. Prior work has found that Schwann cells are eliminated in the distal nerve stump following complete neonatal denervation but not adult denervation [42]. Thus, the loss of Schwann cells may be responsible for the formation of contractures following neonatal but not adult postganglionic brachial plexus injury [6, 43]. The fate and function of Schwann cells and muscle glial cells following neonatal preganglionic denervation is unknown, but preservation of Schwann cells and muscle glial cells accompanying preservation of afferent and sympathetic innervation in preganglionic injury may be a putative mechanism for the protective effect of preganglionic injury against contractures.

Despite not playing a central role in contracture formation, sympathetic neural input regulates critical aspects of skeletal muscle function and maintenance through β-adrenergic signaling [25, 26]. In the current study, we extend prior knowledge of this neuronal pathway by investigating its role in the neonatal growth of normally innervated muscles. Here, we report a sex-specific requirement of sympathetic innervation for developmental muscle growth in male mice, as demonstrated following chemical ablation of sympathetic neurons. Our discovery of this sex dimorphism is complemented by observations of increased muscle growth only in neonatal male mice and rats with β_2_-agonist treatment. Early studies in juvenile rats demonstrated a beneficial effect of clenbuterol treatment in denervated soleus muscles, but did not identify sex dimorphisms as only male rats were tested [44]. Although we did not observe a beneficial effect in denervated brachialis muscles, the neonatal time period assessed in the present study differs from prior investigations on denervation atrophy with clenbuterol. While the misuse of β_2_-agonists (including clenbuterol) as an image– and performance-enhancement drug has been documented within the athletic community [45], the mechanisms by which these sympathomimetic agents confer muscle growth and reduce atrophy are ill-defined [46]. Recent studies have identified increased myonuclear accretion in fast-twitch muscles of adult male rats [47], as well as elevated cAMP/PKA and Akt2 signaling in resistance-trained young male adults [48] as putative molecular adaptations. Whether these mechanisms are conserved during the neonatal stage of muscle development must be further investigated to establish concrete insights into β_2_-agonists as a potential therapy for reversing muscle wasting in younger patients.

Besides anabolic actions, β_2_-agonists exhibit thermogenic properties that may prove valuable for combating obesity and improving metabolism [49]. Specifically, clenbuterol’s desired effect on weight loss has led to a recent rise in its misuse by teenage females [50, 51]. Recent advances reported that clenbuterol improves glucose homeostasis by activating and coupling skeletal muscle β_2_-adrenergic receptors in skeletal muscle G_s_ protein [52]. Furthermore, clenbuterol was self-reported by female users to be associated with minimal androgenic adverse effects, in comparison to anabolic steroids [53]. It is thus not inconceivable that clenbuterol preferentially targets adipose tissues and/or facilitates muscle cell metabolic reprogramming, rather than protein anabolism in female users. However, clenbuterol also attenuated bone growth in our neonatal female mice, which corroborates prior findings of impaired bone mineral density and increased bone resorption in adult male rats [54], and argues against its use as a potential antidiabetic therapy for young female patients. Taken together, our collective results identify a promising molecular target for the treatment of muscle wasting disorders in young male patients, while concomitantly cautioning against the use of β_2_-agonists in young female patients. Importantly, these findings complement our recent discovery that myostatin inhibition facilitates greater growth of innervated muscles and preferentially rescues denervated muscle atrophy in neonatal female mice [15], thereby offering unique therapeutic opportunities for both young female and male patients with muscle diseases. Moreover, our findings underscore the importance of accounting for sex as a biological variable (SABV) [55] when dissecting the molecular mechanisms of muscle growth and the pathophysiology of muscle atrophy [15].

Critically, the divergence of increased muscle CSA in normally innervated muscles and exacerbated sarcomere overstretch in denervated muscles with clenbuterol reveals potentially distinct mechanisms involved in the neonatal development of skeletal muscle width and length. While length is an important facet of muscle development, the underlying mechanism(s) that drive longitudinal muscle growth remains poorly understood [56, 57]. This dearth in our current understanding severely impedes clinical efforts to treat myobrevopathies – disorders stemming from muscle shortness [58]. By uncoupling muscle width and length, we discover that the neurological environment plays a vital role in differentially regulating cross-sectional and longitudinal muscle growth. Here, our results reveal that altered β_2_-adrenergic signaling in the presence of neural input impacted CSA, but alterations in this pathway in the complete absence of neuronal innervation impacted sarcomere length in both neonatal male mice and rats. In total, β-adrenergic signaling increases the ratio of CSA to length, in innervated muscle by increasing CSA, and in denervated muscle by decreasing length. This divergent role of the neurological environment is supported by our prior findings whereby neonatal pharmacologic proteasome inhibition prevented sarcomere overstretch in denervated muscles but impaired cross-sectional growth of innervated muscles [14]. Additional work is required to fully dissect the underlying mechanisms by which neural input differentially regulates CSA and longitudinal growth in a developing muscle. Moreover, future studies must consider both muscle width and length as distinct but pertinent facets of skeletal muscle growth.

## Conclusion

In conclusion, we discover several key insights that advance our understanding of both contracture pathophysiology and skeletal muscle biology. First, our findings rule out sympathetic muscle innervation and its primary molecular pathway in the modulation of contractures, thereby directing our focus on alternative targets for pharmacologic therapies. Second, our discovery of β_2_-adrenergic agonist-mediated sex dimorphisms in neonatal muscle growth offers potential therapies for treating muscle wasting in young male patients, and highlights the importance of SABV in mechanistic and translational studies. Lastly, the discovery of potentially divergent regulation of muscle width and length advocates for their uncoupling in future works interrogating the mechanistic underpinnings of skeletal muscle development.

## Materials and Methods

### Ethical Statement

This study was performed in strict accordance with recommendations in the Guide for the Care and Use of Laboratory Animals of the National Institutes of Health. All rodents were handled according to approved institutional animal care and use committee (IACUC) protocols (#2020-0067) of the Cincinnati Children’s Hospital Medical Center, and every effort was made to minimize suffering.

### Rodents

The following two species were used in this study: wildtype mice (Charles River; CD-1® IGS mouse, strain code 022), and wildtype rats (Charles River; Sprague-Dawley® rat, strain code 400).

### Neonatal Brachial Plexus Injury (NBPI)

Two models of NBPI were surgically created in postnatal day (P)5 rodents under isoflurane anesthesia. Preganglionic NBPI was generated in P5 wildtype female and male mice by unilateral intra-foraminal transection of the C5 – C7 dorsal and ventral rootlets, which preserves afferent and sympathetic innervation to the muscle, and does not induce contractures (**Fig. 3A**) [8]. In contrast, postganglionic NBPI was achieved in P5 wildtype female and male mice and rats by unilateral extraforaminal excision of the C5 – T1 nerve roots, which causes complete efferent, afferent, and sympathetic denervation and induces contractures (**Fig. 5A**) [8]. The different nerve root levels selected for preganglionic (C5 – C7) and postganglionic (C5 – T1) surgeries were chosen due to mortality encountered with C5 – T1 preganglionic injury, as well as potential for spontaneous recovery of C5 – C7 postganglionic injury, complicating the contracture phenotype. Both surgeries denervate the elbow flexor muscles, which are used as the model for contracture investigation in this study. To ensure permanent deficits and prevent potential confounding consequences of reinnervation, deficits in motor function were validated postoperatively and prior to sacrifice. Rodents that displayed preserved or recovered movement in the denervated limb were excluded from the study. Based on this criteria, 9 mice were excluded from the study following preganglionic NBPI, and 1 rat was excluded from the study following postganglionic NBPI.

### Deafferentation and Deefferentation

Neonatal combined deafferentation and deefferentation were surgically created in P5 wildtype female and male rats under isoflurane anesthesia. This mode of denervation was achieved through unilateral dorsal root ganglion excision combined with ventral rootlet transection of the ipsilateral C5 – C7 nerve roots, proximal to the sympathetic chain, thus preserving only sympathetic innervation to the muscle. Similar to the NBPI surgeries, deficits in motor function and active movement in the denervated limb were assessed postoperatively and before sacrifice. Based on the criteria described above, no rats were excluded from the study.

### Cervical Sympathectomy

Neonatal cervical sympathectomy was surgically created in P5 wildtype female and male rats under isoflurane anesthesia. Briefly, subcutaneous tissues were dissected to expose the omohyoid muscle, which was retraced caudally to the level of the clavicle. Sympathectomy was generated by unilateral excision of the superior cervical ganglion located at the level of carotid bifurcation, along with the middle and inferior/stellate ganglia located between the subclavian artery and aortic arch. Rats were assessed post-surgery to confirm ipsilateral eye ptosis (Horner’s syndrome). Non-sympathectomized, contralateral forelimbs served as controls.

### Chemical Sympathectomy

To chemically ablate sympathetic neurons following preganglionic surgery, mice were treated with the neurotoxin 6-hydroxydopamine (6-OHDA). Starting at one day post-NBPI, mice were initially administered 100mg/kg 6-OHDA (Sigma-Aldrich #162657) for one continuous week, followed by once weekly treatments thereafter through subcutaneous injections [59]. Saline was administered in a separate litter of mice as controls. In a separate experiment, two independent litters of mice were administered either 6-OHDA or saline in the absence of preganglionic surgery to assess the effect of chemical sympathectomy in normally innervated muscles.

### Beta_2_-Agonist

To stimulate β-adrenergic activity after post-ganglionic surgery, mice and rats were treated with the β_2_-agonist, clenbuterol (Sigma-Aldrich #C5423). Following post-ganglionic NBPI at P5, mice and rats were administered 3 mg/kg and 5 mg/kg of clenbuterol daily through subcutaneous injections, respectively. Dulbecco’s phosphate-buffered saline (DPBS) served as control in separate litters of mice and rats.

### Animal Husbandry

Post-surgery (NBPI and cervical sympathectomy), all rodents were returned to their respective mums and housed in standard cages on a 1:1 light/dark cycle, with nutrition and activity *ad libitum*. All control and experimental rodents in this study were randomized by litter, with all drug treatments (6-OHDA and clenbuterol) administered at noon and in the respective cages. To account for behavioral changes reported with both drugs [60, 61], rodents were monitored for signs of aggression and distress. Daily body weights were also recorded for all rodents to monitor the toxicity of the drug treatments.

### Assessment of Contractures

At P33 (4 weeks post-NBPI, cervical sympathectomy, and chemical sympathectomy), all rodents were euthanized by CO_2_ asphyxiation. Images of bilateral elbow and shoulder range of motion (ROM) were captured at maximum passive extension and external rotation, respectively, with a validated digital photography technique [13–15, 22]. Elbow flexion and shoulder internal rotation contractures were subsequently calculated by measuring and comparing elbow and shoulder ROM in AxioVision (Zeiss). Both digital photography and image analysis were performed with blinding to treatment groups. Representative images of forelimbs shown within **Figs. 1B, 2B, 3C, 5C, 6C, 7B** have been processed to reflect comparable levels of sharpness, brightness, and contrast for illustrative purposes. No image manipulation was performed prior to measurements.

### Tissue Collection/Preparation

Following image photography, the musculocutaneous nerve (MCN) in bilateral forelimbs were dissected and fixed in 4% paraformaldehyde (PFA) for 5 h, and stored in PBS at 4°C for immunohistochemical staining. The remaining forelimbs were positioned on cork at 90° elbow flexion, imaged by digital x-ray for humerus length and symmetry of joint positioning between bilateral limbs, and fixed in 10% formalin for 48 h. Bilateral brachialis muscles were subsequently removed, soaked in 25% Lugol solution (Sigma-Aldrich #32922) overnight, and processed for micro-computed tomography (MicroCT) to assess whole muscle size [13–15, 22]. Post scanning, muscles were recovered in PBS overnight at 4°C, and digested in 15% sulfuric acid for 30 min the following day. Muscle bundles were then isolated and imaged for sarcomeres via differential interference contrast (DIC) microscopy at 40x on a Nikon Ti-E SpectraX widefield microscope [13–15, 22].

### *Ex-Vivo* High Resolution Studies

MicroCT was performed using a Siemens Inveon PET/SPECT/CT scanner (Siemens Medical Solutions) as previously described [15, 22]. Briefly, acquisitions were reconstructed using a Feldkamp algorithm with mouse beam-hardening correction, slight noise reduction, 3D matrix size 1024×1024×1536, according to manufacturer-provided software. Protocol-specific Hounsfield unit (HU) calibration factor was applied. The cone-beam CT parameters were as follows: 360° rotation, 1080 projections, 1300 ms exposure time, 1500 ms settle time, 80 kVp voltage, 500 μA current, and effective pixel size 17.67 μm.

### Morphometrics

Humerus lengths were quantified through the AxioVision program by measuring the distance between the proximal humerus physis to the distal articular surface on digital radiographs [13–15, 22]. Brachialis whole muscle measurements were obtained by processing the MicroCT scans into digital imaging and communications in medicine (DICOM) images with Fiji programs, and subsequently normalizing to humerus length of the corresponding forelimb [13–15, 22]. Specifically, whole muscle cross-sectional area (CSA) was quantified in Segmentation Editor, whereas muscle volume was processed in 3D Viewer. Average brachialis sarcomere length was determined by measuring series of 10 sarcomeres from 6 representative DIC images from each muscle with AxioVision software. The presence of elongated sarcomeres is associated with a fewer number of sarcomeres in series, thereby indicating a reduction in functional length and longitudinal growth of the brachialis, as previously described [13–15]. All morphometric measurements were performed with blinding to treatment groups. For illustrative purposes, raw DICOM files were processed in IMARIS software (Bitplane) to create whole-muscle images presented in **Figs. 4B, 8B, 9B**. Representative images of sarcomeres shown within **Figs. 1E, 2D, 3F, 5F, 6F, 7E** have been cropped to identical sizes, and processed to reflect comparable levels of sharpness, brightness, and contrast for illustrative purposes. No image manipulation was performed prior to measurements.

### Whole-mount Immunofluorescence

Following fixation, MCNs were permeabilized with 1% Triton X-100, and blocked in 10% donkey serum/1% bovine serum albumin (BSA) overnight at 4°C. MCN samples were subsequently incubated for 72 h at 4°C with primary antibodies against Neurofilament Heavy protein (NF-H) (1:2500; Novus #NB300-217) and tyrosine hydroxylase (1:250; Novus #NB300-110). Following another 48 h of incubation at 4°C with secondary Alexa Fluor antibodies (1:100–200; Invitrogen), samples were cleared overnight in Refractive Index Matching Solution (RIMS) buffer containing 80% iohexol (Histodenz™) [62]. MCN samples were finally mounted with ProLong™ Gold Antifade Mountant and stored overnight. Rat MCN whole-mounts were visualized at 10x on a Nikon A1R confocal system, whereas mouse MCN whole-mounts were visualized at 4x and 10x on a Nikon Ti-E SpectraX widefield microscope.

### Statistics

For all data sets, outliers were detected *a priori* by Grubbs’ test and excluded. To compare differences in values between two data sets, all data were tested for normality with the Shapiro–Wilk test. Normally distributed data were subsequently compared with unpaired two-tailed Student’s t-tests, whereas non-normally distributed data were compared with Mann–Whitney U-tests. For data sets with two independent variables (sex and treatment), a two-way analysis of variance (ANOVA) with Bonferroni correction for multiple comparisons was performed [15]. For data sets with three independent variables (sex, treatment, denervation), a three-way ANOVA with repeated measures between forelimbs and Bonferroni correction for multiple comparisons was performed [15]. All data are presented as mean ± SD. The degree of significance between data sets is depicted as follows: *p<0.05, **p<0.01, ***p<0.001, ****p<0.0001. *A priori* power analyses based on prior work determined that 4-6 mice per group were required to detect a 10° difference in contractures and a 0.2 µm difference in sarcomere lengths at 80% power between experimental conditions, or a 50% therapeutic reduction in contracture severity. A total of 10 rats and 74 mice were used in this study. All statistical tests were performed with Prism 8 software (GraphPad).

### Author Contributions

QG and RC conceived and supervised the study; MTR, AT, QG, and RC designed the research and experiments; MTR, AT, KSW, MEE, QG, and RC performed experiments; MTR, AT, KSW, and MEE curated data; MTR, AT, KSW, MEE, QG, and RC analyzed data; QG wrote the manuscript with assistance from all authors.

## Acknowledgements

We thank the following entities within Cincinnati Children’s Hospital Medical Center: the Veterinary Services for surgical assistance, and the Confocal Imaging Core for microscope assistance. We also thank the following individuals within the University of Cincinnati College of Medicine: Sharon Wang from the Preclinical Imaging Core for MicroCT assistance, and Dr. Sarah Kurkowski from the Department of Orthopaedic Surgery for illustrating the Graphical Abstract. This work was supported by grants to RC from the National Institutes of Health (NIH) (R01HD098280-01), as well as funding from the Cincinnati Children’s Hospital Division of Orthopaedic Surgery and Junior Cooperative Society. The respective funding sources were not involved in the study design; in the collection, analysis, and interpretation of data; in the writing of the report; and in the decision to submit the paper for publication.

## Data Availability Statement

All data generated or analyzed during this study are included in the manuscript and supporting files.

## List of Supplementary Figures

**Figure 1 – supplemental figure 1:**
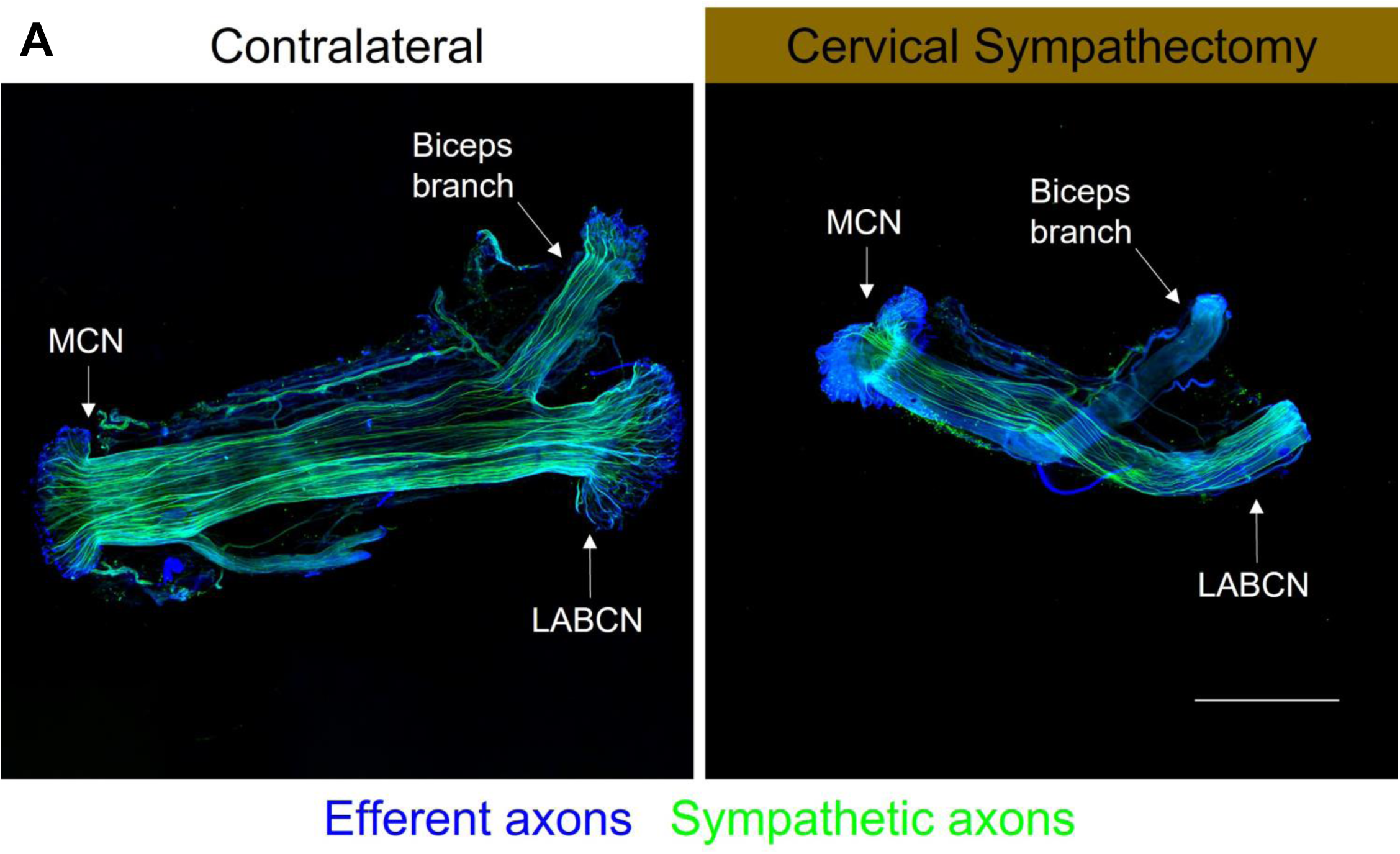
Immunostaining of musculocutaneous nerves following surgical sympathectomy. (**A**) Whole mounts of rat musculocutaneous nerves (MCNs) confirmed that 4 weeks of cervical sympathectomy preserves motor axons but successfully ablates sympathetic axons along the sympathetic chain leading to the biceps. Scale bar: 1000 µm.

**Figure 2 – supplemental figure 1:**
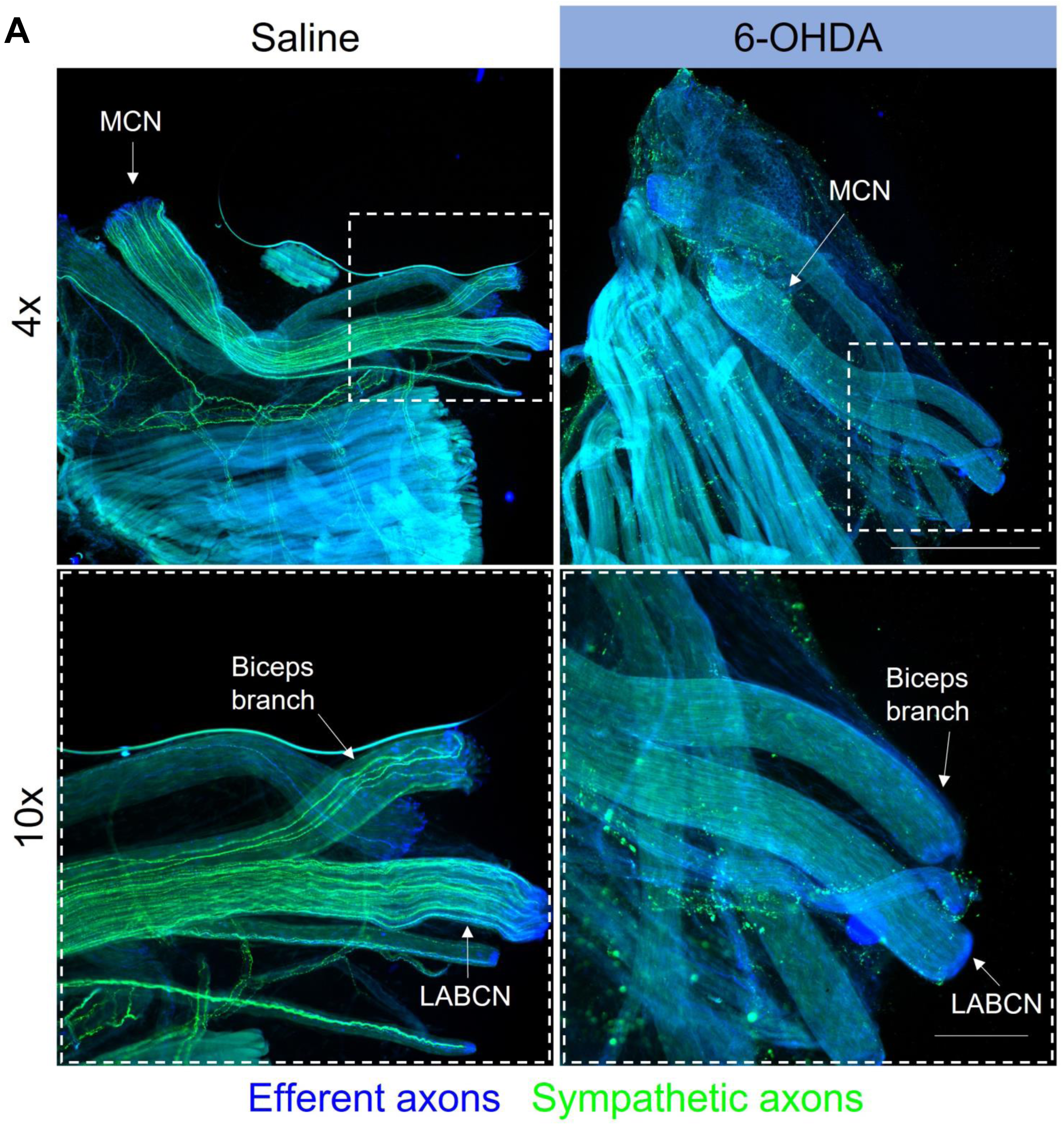
Immunostaining of musculocutaneous nerves following chemical sympathectomy. (**A**) Whole mounts of mouse MCNs verified the presence of motor axons and the absence of sympathetic axons throughout the nerve following 4 weeks of 6-OHDA treatment. The top row depicts images taken at 4x magnification. The dotted boxes represent the corresponding insets depicted on the bottom row, which were taken at 10x magnification. Scale bars: top row – 1000 µm, bottom row (insets) – 250 µm.

**Figure 4 – supplemental figure 1:**
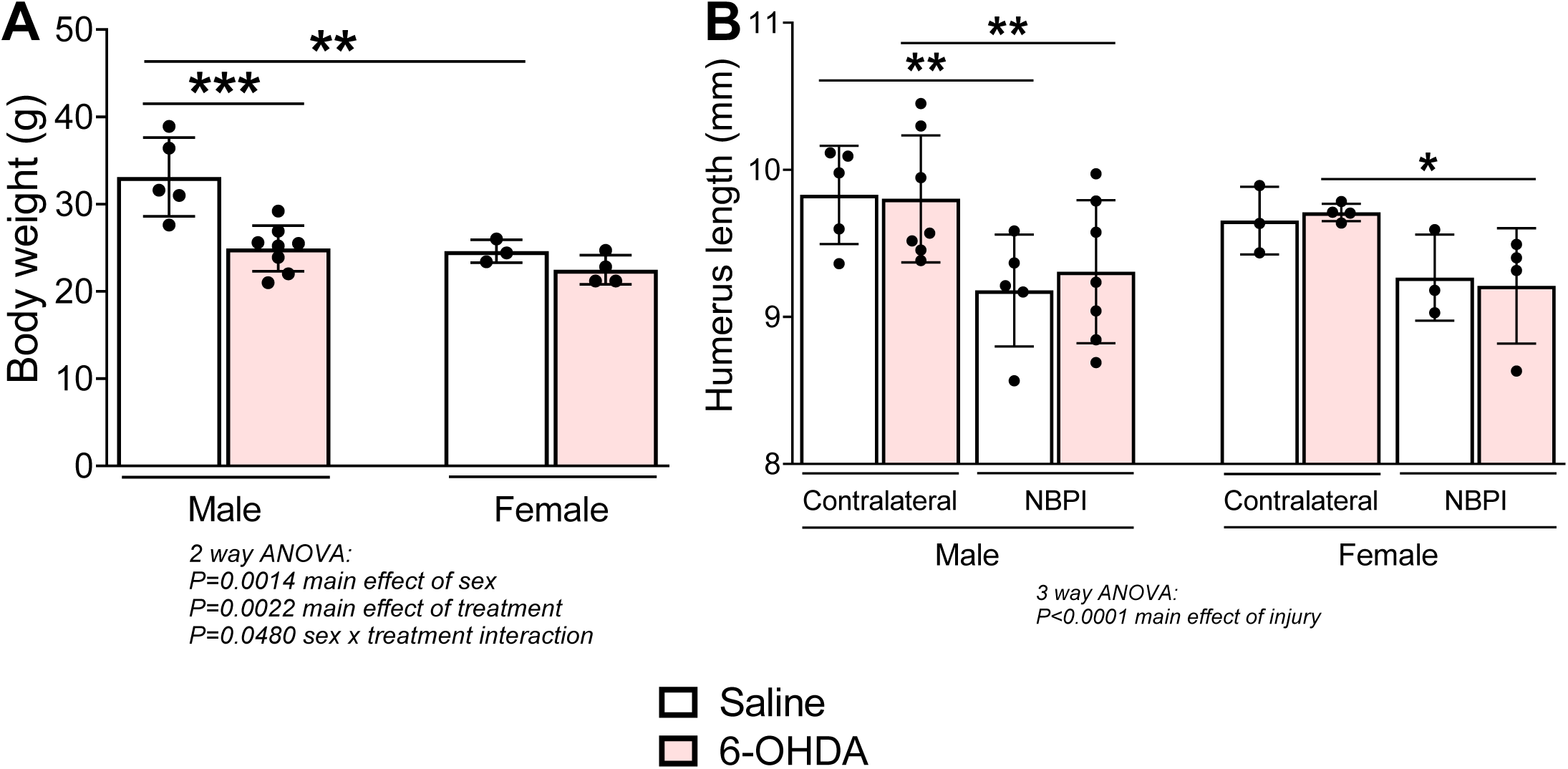
Effect of chemical sympathectomy on neonatal growth. (**A**) 6-OHDA treatment impeded terminal body weight in male mice, (**B**) but did not alter humerus lengths in control and denervated forelimbs of both male and female mice. Data are represented as mean ± SD, n = 3-8 independent mice per group. Statistical analyses: (**A**) two-way ANOVA for sex and treatment with a Bonferroni correction for multiple comparisons, (**B**) three-way analysis of variance (ANOVA) for sex, treatment, and denervation (repeated measures between forelimbs) with a Bonferroni correction for multiple comparisons. *p<0.05, **p<0.01, ***p<0.001.

**Figure 8 – supplemental figure 1:**
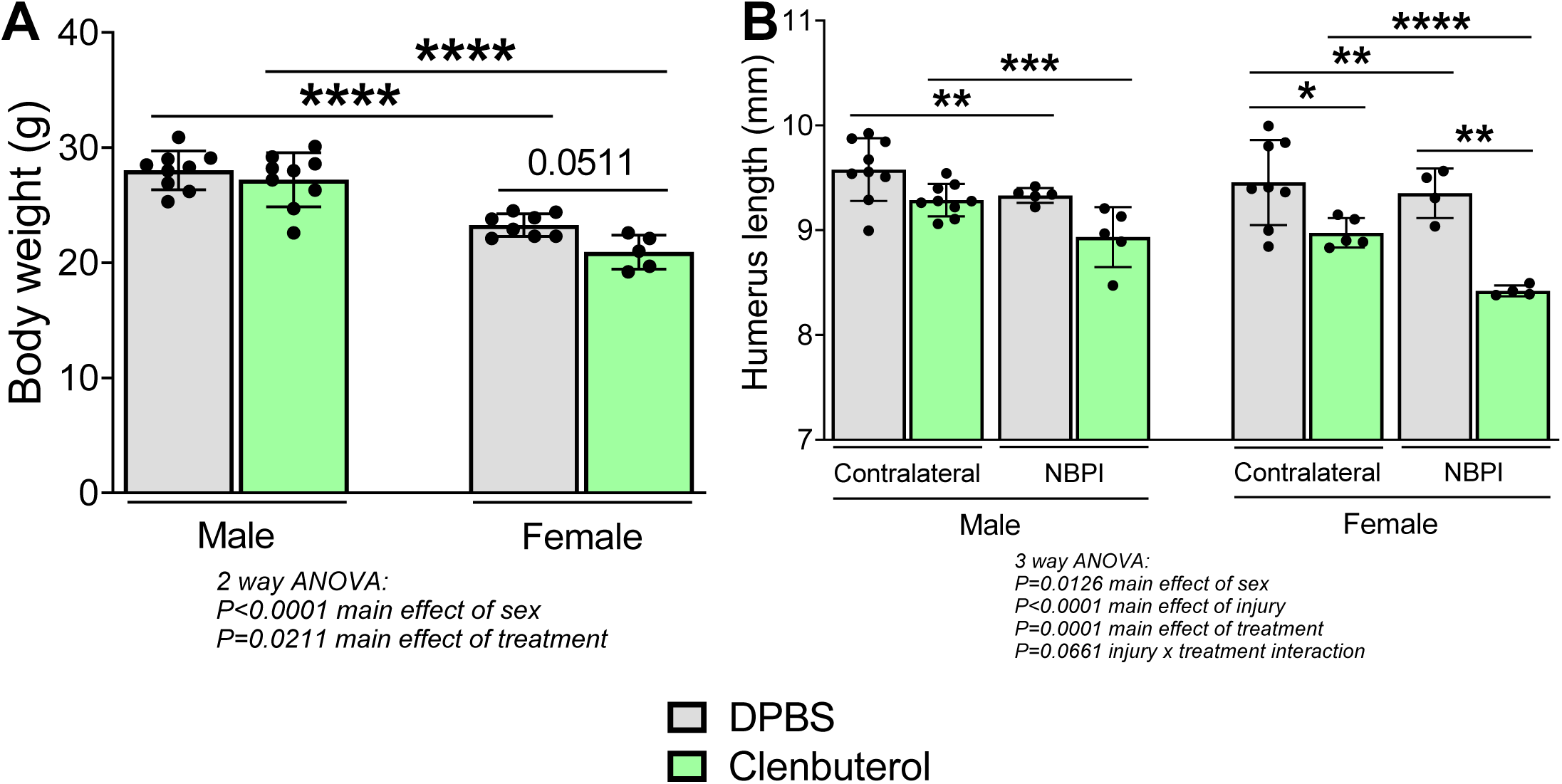
Effect of β-adrenergic stimulation on neonatal growth. Clenbuterol treatment did not significantly alter body weight (**A**) but reduced (**B**) humerus lengths in control and denervated forelimbs of female mice. Data are represented as mean ± SD, n = 4-9 independent mice per group. Statistical analyses: (**A**) two-way ANOVA for sex and treatment with a Bonferroni correction for multiple comparisons, (**B**) three-way analysis of variance (ANOVA) for sex, treatment, and denervation (repeated measures between forelimbs) with a Bonferroni correction for multiple comparisons. *p<0.05, **p<0.01, ***p<0.001, ****p<0.0001.

